# AlphaFold3 at CASP16

**DOI:** 10.1101/2025.04.10.648174

**Authors:** Arne Elofsson

**Affiliations:** Dep of Biochemistry and Biophysics and Science for Life Laboratory, Stockholm University

## Abstract

The CASP16 experiment provided the first benchmark for AlphaFold3. In contrast to AlphaFold2 and other methods, AlphaFold3 can also predict the structure of non-protein molecules. In this study, we assess the performance of AlphaFold3 using both automatic server submissions and manual predictions from the Elofsson group. All predictions were generated via the AF3 web server, with manual interventions applied to large targets. Our analysis shows that AF3 performs comparably to top predictors for proteins and complexes, with average GDT_TS and DockQ scores indicating high model quality. Manual curation did not significantly improve results over default AF3 server submissions. Compared to AlphaFold2-based methods, we found that AF3 performs slightly better for protein complexes. Still, when massive sampling is applied to AlphaFold2, the difference disappears, using standard measurements. The performance of the AlphaFold3 server is comparable to the best methods when only taking the top-ranked predictions into account, but slightly behind when examining the best out of the five submitted models. Further, there are many targets where one method makes a good prediction while another top-ranked method fails, indicating that a venue for progress could be to develop better methods for identifying the best models. In the official ranking from CASP, AlphaFold3 performs better than AlphaFold2 for easier targets, but not for harder targets. Finally, RNA predictions remain difficult, and the accuracy of stoichiometry predictions is limited, especially for heteromeric targets. Overall, AlphaFold3 provides an easy-to-use method that offers close to state-of-the-art predictions.

## Introduction

Protein structure prediction has revolutionized in the last decade, starting with rediscovering DCA^1,2^ and other methods to predict contact from multiple sequence alignments. It was shown that these methods could accurately predict the structure of proteins given sufficiently deep multiple sequence alignments^3–5^. These methods were then improved by using various machine learning methods to post-process these predictions^3,6–8^. From the perspective of CASP, the progress could first be seen at CASP13 when AlphaFold(v1)^9^ performed better than other methods for challenging protein targets^10^ with the introduction of AlphaFold2.0^11^.

In addition to predicting the structure of individual proteins, contact prediction methods can be used to predict the structure of complexes^12^. So-called Fold and Dock approaches showed progress in CASP14 using machine learning enhanced contact predictions^13^. With the release of AlphaFold2, these methods suddenly could predict the structure of many complexes^14^. CASP15 showed that methods based on AlphaFold2 could predict the structure of these complexes accurately^15^. One challenge that seemed to remain was the prediction of AntiBodies/NanoBodies to Antigens, but increased sampling helped in some cases^16,17^.

However, proteins are not the only molecules in a cell; nucleotides, lipids, ions, and small molecules are also present. In 2024, several tools to predict the interactions between all these types of molecules were released ^18,19^, and CASP16 was the first opportunity to benchmark these in a completely blind fashion. At the time of the CASP16 submissions, AlphaFold3 was only available as a web server, limiting its use. The main limitation was that only a few standardized protein ligands could be used.

For CASP16, I submitted manual predictions as group Elofsson (#241) and server predictions as AF3-server (#304). All predictions were made using the AlphaFold3 server^1^. The five models from one target were submitted for the server predictions. In contrast, more models were generated, and good models with some structural variability were submitted for the manual predictions. For some models, additional changes were made; see methods. Here, I will describe the results briefly.

## Methods

We set up to use the AlphaFold3 server for all predictions in CASP16, as it was impossible to install it locally at the time of CASP16. We had to do manual interventions to make it work for all targets, but we tried to use it as automatically as possible. The manual interventions were necessary for all ligand targets with unknown stoichiometry and targets bigger than 5000 residues. Details for how this was done are described below.

In addition to the automatic predictions as AF3-server (#304), we also submitted manual predictions as the Elofsson group (#241). The only difference for the Elofsson predictions was that we generated additional models by running AlphaFold3 with different random seeds. Then, we used the internal AlphaFold3 submission and some manual inspections to select the models to submit. As can be seen from the results below, the manual intervention did not provide any significant improvement, so it will not be discussed in detail here.

### Predictions

#### Big targets

Some targets (e.g., H1227, H1257) were too big to run on the server (the maximum is 5000 residues/ nucleotides). For these targets, we used the strategy from MolPC^14^, i.e., building different types of overlapping fragments, superposing the shared parts, and generating complete models. The cutting was not fully automated for the AlphaFold3 predictions, as we tried to select the best place to provide the cut. However, for AlphaFold3 we only used one cut and submitted the top five models, while for Elofsson we tried several versions. No refinements were done in any cases.

#### RNA

For multichain RNA molecules, it was noted that many models contained significant overlap. These can easily be detected from the scores provided by the AlphaFold3 server, as the flag “has_clash” is set and the “ranking_score” is negative. We excluded these for submission whenever possible. However, this necessitated additional submissions to the server for these targets until five non-clashing models were obtained.

#### Ligand predictions

For ligand targets, we (1) generated a structural model from the smiles using online tools and (2) ran the AlphaFold3 Server with the available ligand most resembling the ligand that should be docked. Finally, the CASP ligand was superposed on the AlphaFold3, generating a new PDB/Ligand file. Unfortunately, we submitted the ligands in PDB format without receiving any errors, so they were not officially evaluated by the prediction center (we provided them in the correct format later, but we assume that was too late).

#### Ensemble targets

Different strategies were used for different ensemble targets.

- R1203, T1214, R0283 Srandard predictions using AlphaFold3
- T1294, T1200/T1300, M1239, T1249, M1228, T2249, R1253. Generated many models and submitted ensembles of them after clustering.
- R1260 (solvation shell): For AF3, an ensemble of models was generated and hydrated using standard protocols. Elofsson took the best models and ran a 47-ns-long MD simulation.

### Unknown stoichiometry

For targets with unknown stoichiometry, we examined the possibility of predicting stoichiometry by generating models with possible stoichiometries. For computational reasons, it was not possible to generate all possible stoichiometries. Therefore, we used some manual intervention to select which ones to try. We usually ran monomers to hexamers for a start. Because we did not try larger complexes, this caused problems for some targets, including R1253, an octamer, and therefore, failed to predict it accurately. After the initial screen, we used the ranking confidence and selected the five best scores for AF3-submissions. At the same time, for the Elofsson submissions, we submitted a more varied set of predictions, taking manual inspection of the PAE maps into account.

Further, this approach was impossible for targets containing multiple chains, i.e., M and H targets, as the number of combinations was too big. We only used a limited set of combinations for these targets for the AF3 Server. We individually added copies of the individual chains, evaluating the PAE maps for the Elofsson predictions.

### Evaluations

CASP uses many different measures to evaluate the accuracy of a model. All these methods provide other insights. However, most of these measures are highly correlated, and a detailed analysis of all predictions will be presented elsewhere in this issue. Therefore, we will only show results of GDT_TS for monomer and DockQ for complexes. We also present data about the ranking of the methods using the Z-score analysis from the prediction center. All data is downloaded from the prediction center. All analysis scripts are freely available from https://gitlab.com/arneelof/CASP16-predictions. Except for stoichiometry analysis, we only analysed Round 1 models. We did not make any changes for Round 2; the only change between Round 1 and Round 2 was a change of stoichiometry if necessary. In all pairwise comparisons, only the targets by both methods were included.

Only one result for H1265, which appears four times on the prediction centers website, was included: H1265v1. H1265v2 and H1265v3 were ignored.

## Results

In this paper, we analyze the performance of the AlphaFold3 server in CASP16. It should be remembered that all (manual) groups had access to the AF3-server. Therefore, this can be seen as a baseline for all predictions. We primarily focused on comparing the performance of AlphaFold3 in general to other methods and did not focus on the prediction of individual targets. First, we compared the AF3-server predictions with our manual predictions, which were based on running the server more extensively and manually selecting the best model. Secondly, for proteins, we compared the predictions of AlphaFold3 with a baseline AlphaFold2 method, Colabfold_baseline. Finally, we compared the AlphaFold3 models with the best methods according to default scoring on the prediction center website. Comparisons are made separately for pure protein targets and targets that contain nucleic acids.

What is essential to consider is that this paper is not focused on replicating the excellent work of the assessors. Therefore, we use one standard measure, GDT_TS, for the quality of a single domain, and another, DockQ, for the quality of complexes. Indeed, a more detailed analysis could provide additional insights, but these measures are sufficient to detect significant differences between merit methods. If the differences are slight, the gap between the methods is minor. Further, this is not a detailed analysis of individual predictions since our predictions were made with minimal manual intervention. Therefore, we cannot always ascertain why one method performs better than another for a specific target; it often comes down to coincidences. As shown below, a few targets frequently cause one method to appear superior. Even if a difference is statistically significant, it is of minimal importance for the average user if caused by one or a few targets, as these might be coincidences. Therefore, we try to ignore these and only discuss systematic differences between methods.

### Proteins

All models were predicted using the AlphaFold3 server’s default settings. However, if the target was bigger than 5000 residues, it had to be cut into pieces. In these cases, manual decisions were made on where to cut it. The pieces were then merged into a larger protein by superposing the overlapping, as in MolPC^14^. One example of this methodology can be seen in Figure 1.

**Figure 1:**
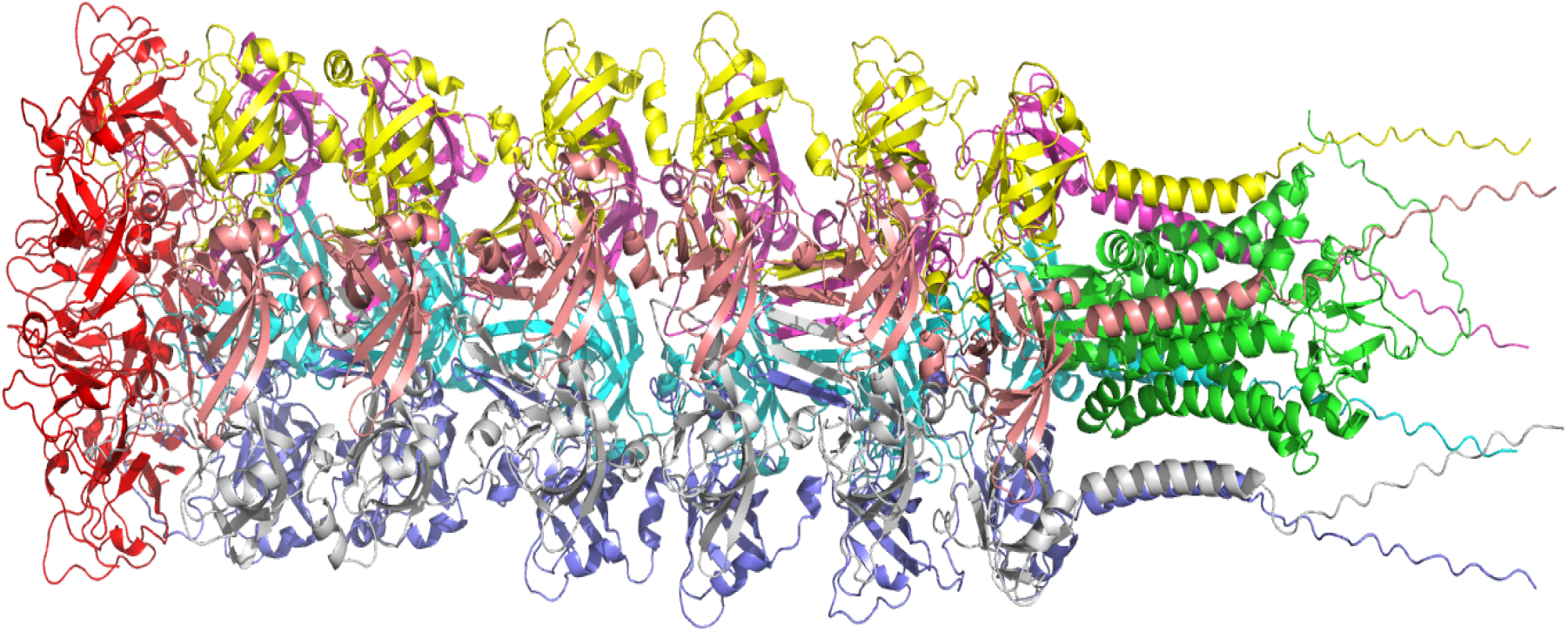
Modeling a large target, H1227, where the left domain (colored in red) could not be modeled directly by AlphaFold3 as the total complex was more than 5000 residues. Instead, it was modeled separately, along with the terminal part of the larger complex. The overlapping domain was superimposed, and the red domain was finally added to the model before submission.

#### Elofsson vs AF3

We first explored whether our manual interventions, submitted under the group Elofsson, were effective. Figure 2 shows that the difference in model quality between AF3-server and Elofsson is slight for the individual domains. When we examine the average GDT_TS scores, AF3 shows slightly better performance. However, this difference originates from two domains: T1212-D1 and T1257-D1. When considering the best of the five predictions, this difference disappears, indicating that, in these cases, AF3-server happened to select a better first-ranked model. Interestingly, it is also worth noting that all but four domains are predicted with high quality, as indicated by a GDT_TS score over 80. The average GDT_TS score is about 88, close to experimental quality.

**Figure 2:**
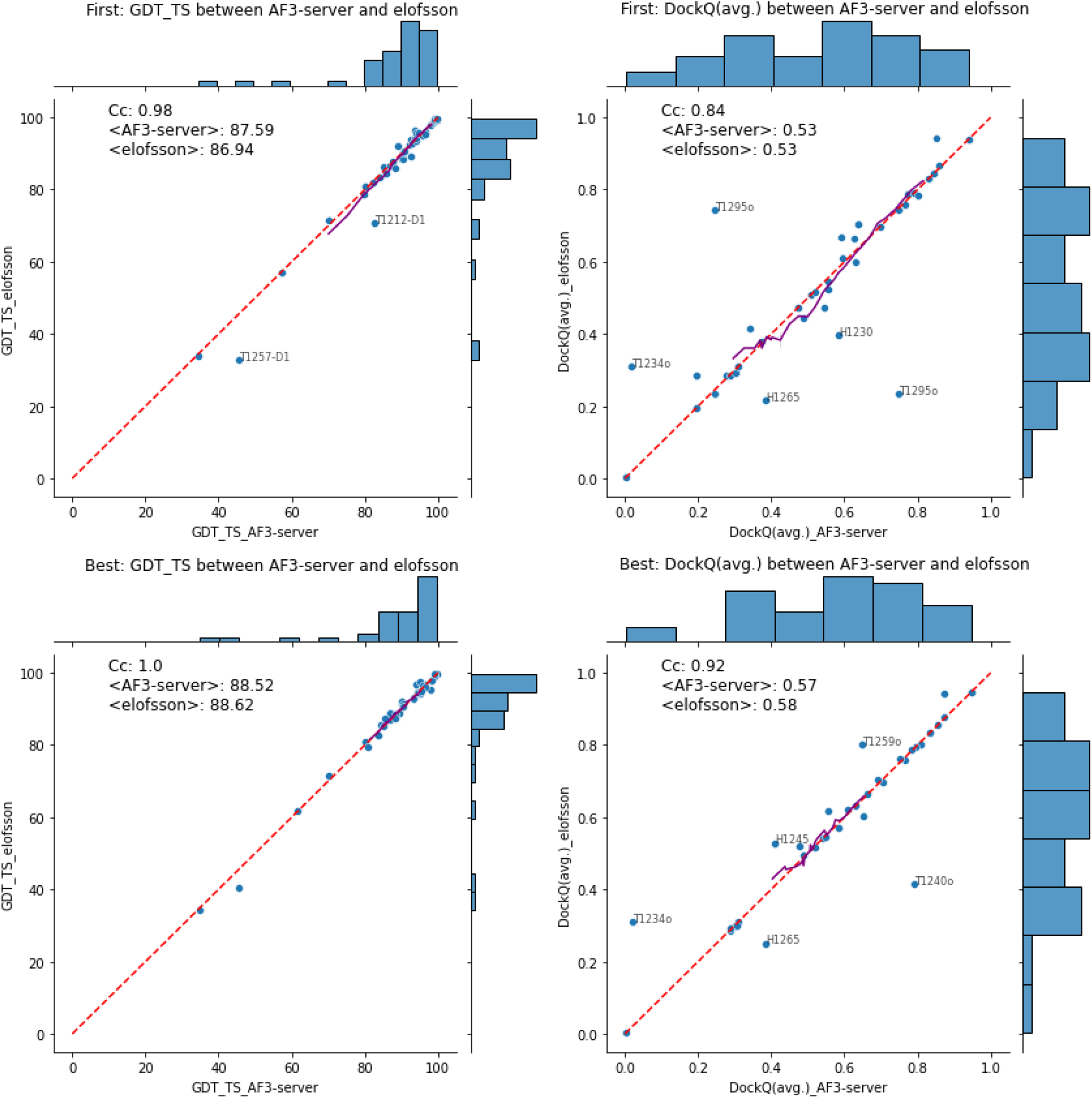
Comparison of AF3-server and Elofsson predictions using GDT_TS and DockQ. The top row compares the highest-ranked prediction, while the second row compares the best prediction out of five. On the left, the comparisons are made at the domain level using GDT_TS, and on the right, comparisons of complexes using DockQ are shown. Any target with a difference greater than 10% of the maximum value for one of the methods is noted. A rolling average over ten points is shown in purple.

However, the correlation for complexes is significantly lower. Again, there is no significant difference between AF3-server and Elofsson, nor when we compare the first-ranked or the best of the submitted models. Most complexes are predicted with acceptable quality, achieving an average DockQ score of 0.53 above the acceptable threshold of 0.23. However, it’s important to note that we are only analyzing average DockQ scores, and many of the complexes have multiple interfaces, so some interfaces may still have low quality.

#### AlphaFold3 vs AlphaFold2

Next, we compared the performance of AlphaFold3 to that of its predecessor, AlphaFold2, see Figure 3. Several versions of what we believe are pure AlphaFold2 predictions are available, including multiple iterations of ColabFold and MassiveFold. Here, we only compared AF3-server with colabfold_baseline and MassiveFold to see if the increased sampling used in MassiveFold affected the performance. AlphaFold3 has a slight edge among the top-ranked domains, with an average GDT_TS of 87.1 compared to ∼85.5 for the AlphaFold2 methods. The difference can be attributed to a few targets where it performed better, notably not the same targets for colabfold_baseline or MassiveFold. The difference fades when examining the best out of five models. Notably, even when studying the best models, there are two domains where AlphaFold3 outperforms the previous version, with one domain showing better results for AlphaFold2.

**Figure 3:**
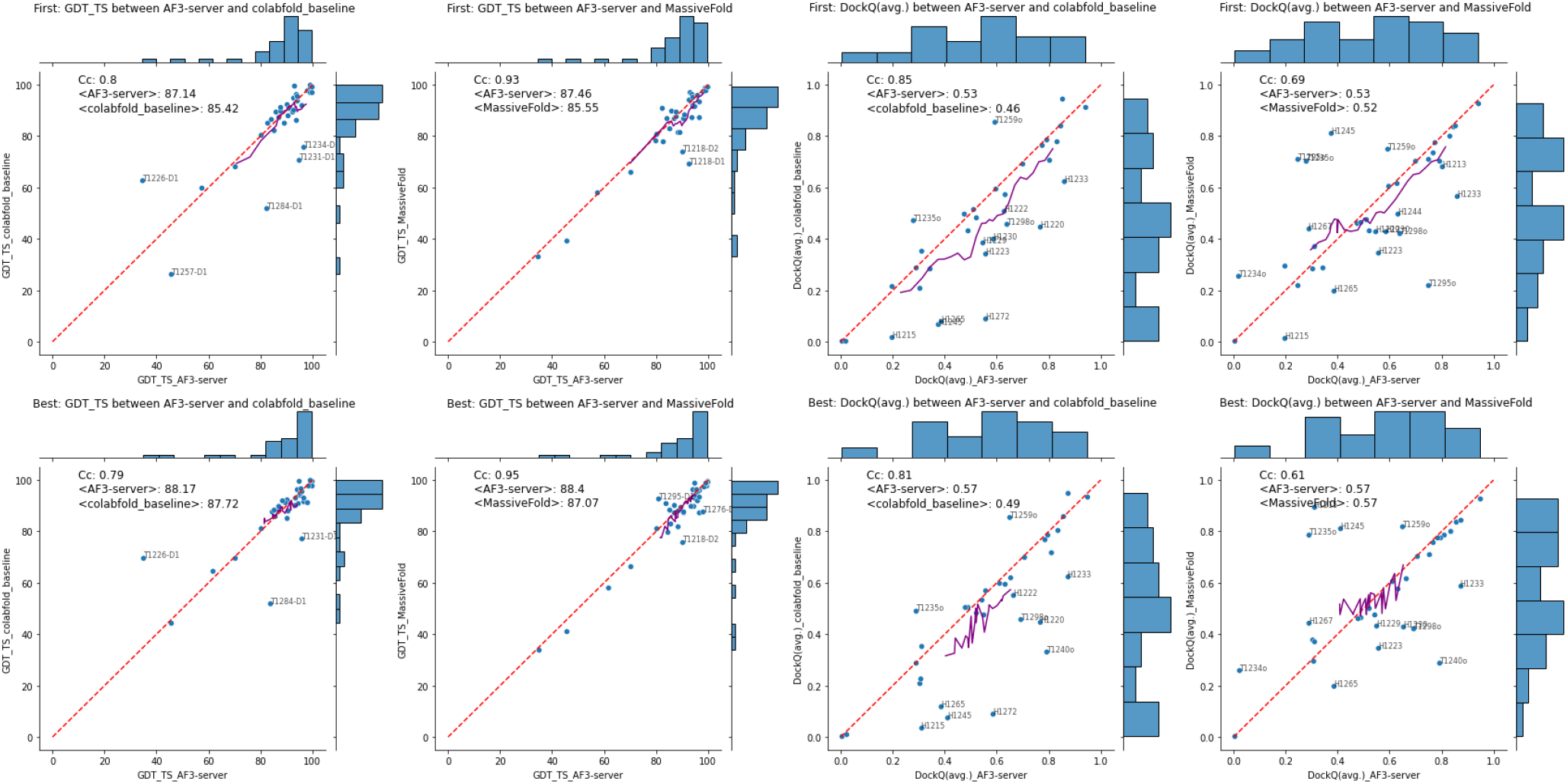
Comparison of AF3-server with MassiveFold and colabfold_baseline predictions using GDT_TS and DockQ. The top row compares the highest-ranked prediction, while the second row compares the best prediction out of five. The comparison is made at the domain level using GDT_TS in the left two columns, and in the right two columns, the comparison is made for complexes using DockQ. Any target with a difference greater than 10% of the maximum value for one of the methods is noted. A rolling average over ten points is shown in purple.

Regarding complexes, AlphaFold3 demonstrates an advantage over colabfold_baseline, achieving an average score of 0.53 compared to AlphaFold2’s 0.46 for first-ranked models and 0.57 versus 0.49 for the best models. Approximately ten AlphaFold3 models are better than corresponding models from colabfold_baseline. However, MassiveFold produces models similar to AlphaFold3 in most cases, indicating that increased sampling enables AlphaFold2 to achieve performance comparable to AlphaFold3.

#### Comparison with top-ranked predictors

Next, we compared AlphaFold3’s performance to that of the top-ranked predictors. It can always be discussed which group performed best overall. We used the prediction center’s default setting to select the top groups, resulting in YangServer for domains and KiharaLab for complexes.

Our analysis showed relatively small performance differences among the top-ranked models for domains and complexes. Still, these differences are more considerable when comparing the best models, as illustrated in Figure 4. The difference can be explained entirely by two targets (T12561-D1 and T1226-D1) for the domains. Notably, when evaluating the complexes, several models from AlphaFold3 outperformed KiharaLab and vice versa. For most models, although the top-ranked AlphaFold3 models perform better, this is no longer the case when considering the best out of the five models.

**Figure 4:**
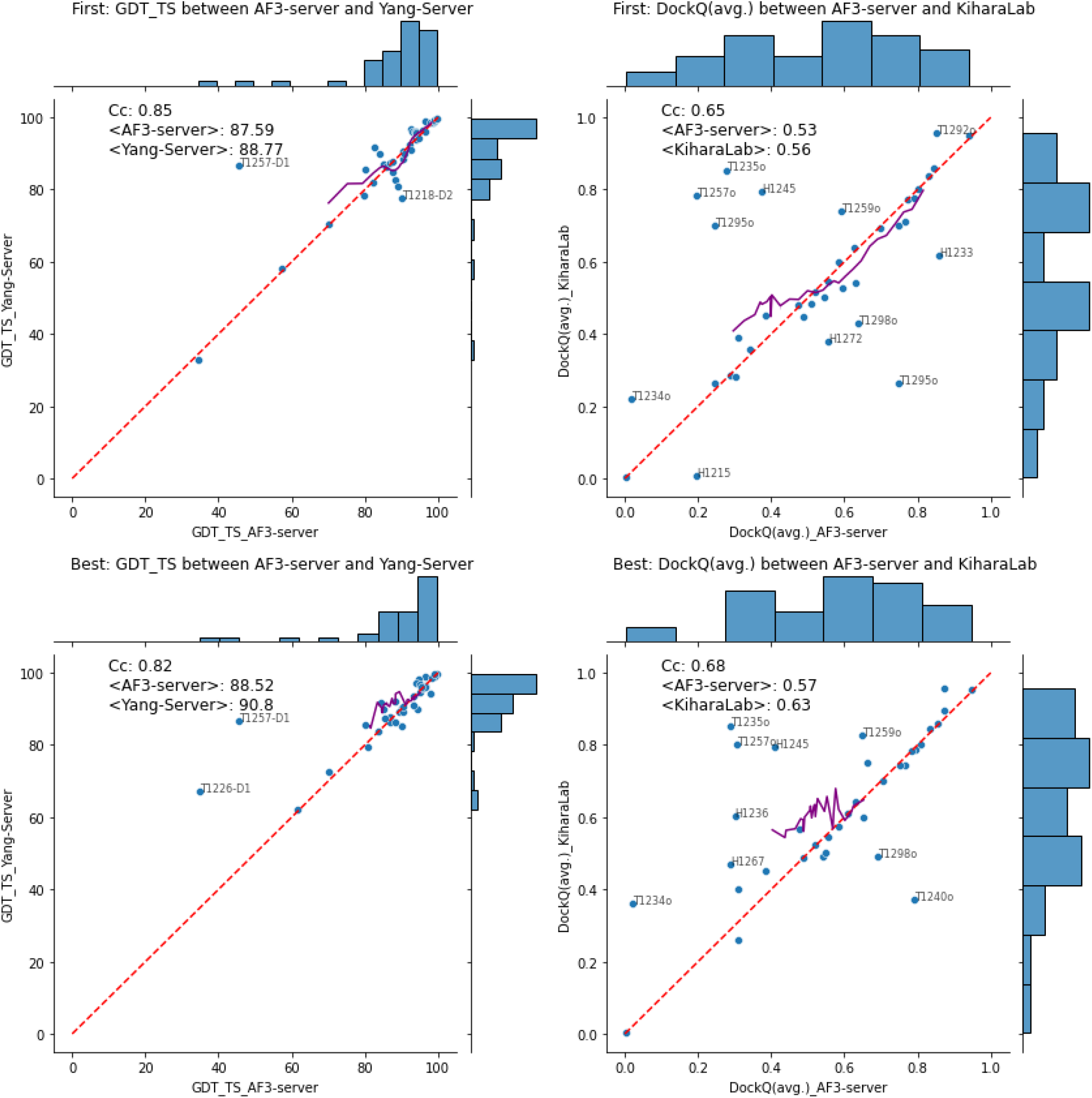
Comparison of AF3-server and top-performing predictors, YangServer for domains and KiharaLab for complexes. The top row compares the highest-ranked prediction, while the second row compares the best prediction out of five. On the left, the comparison is made at the domain level using GDT_TS, and on the right, the comparison of complexes employs DockQ. Any target with a difference greater than 10% of the maximum value for one of the methods is noted. A rolling average over ten points is shown in purple.

Our findings showed that YangServer and KiharaLab generated better “best” out of five models, but not better top-ranked models. The larger improvement when analyzing the best model suggests that the best methods show larger variation when choosing the five models to submit. However, these methods also fail to identify the best submitted models.

#### First vs Best

In Figure 5, we compare the performance of the top-ranked model with the best prediction among all AF3-server predictions. Submitting five models improved the performance for only two complexes: T1240 and T1295. There was no significant improvement for all other models. This shows, again, that for hard prediction cases, generating multiple models can help, but AlphaFold3 can not always detect the best model.

**Figure 5:**
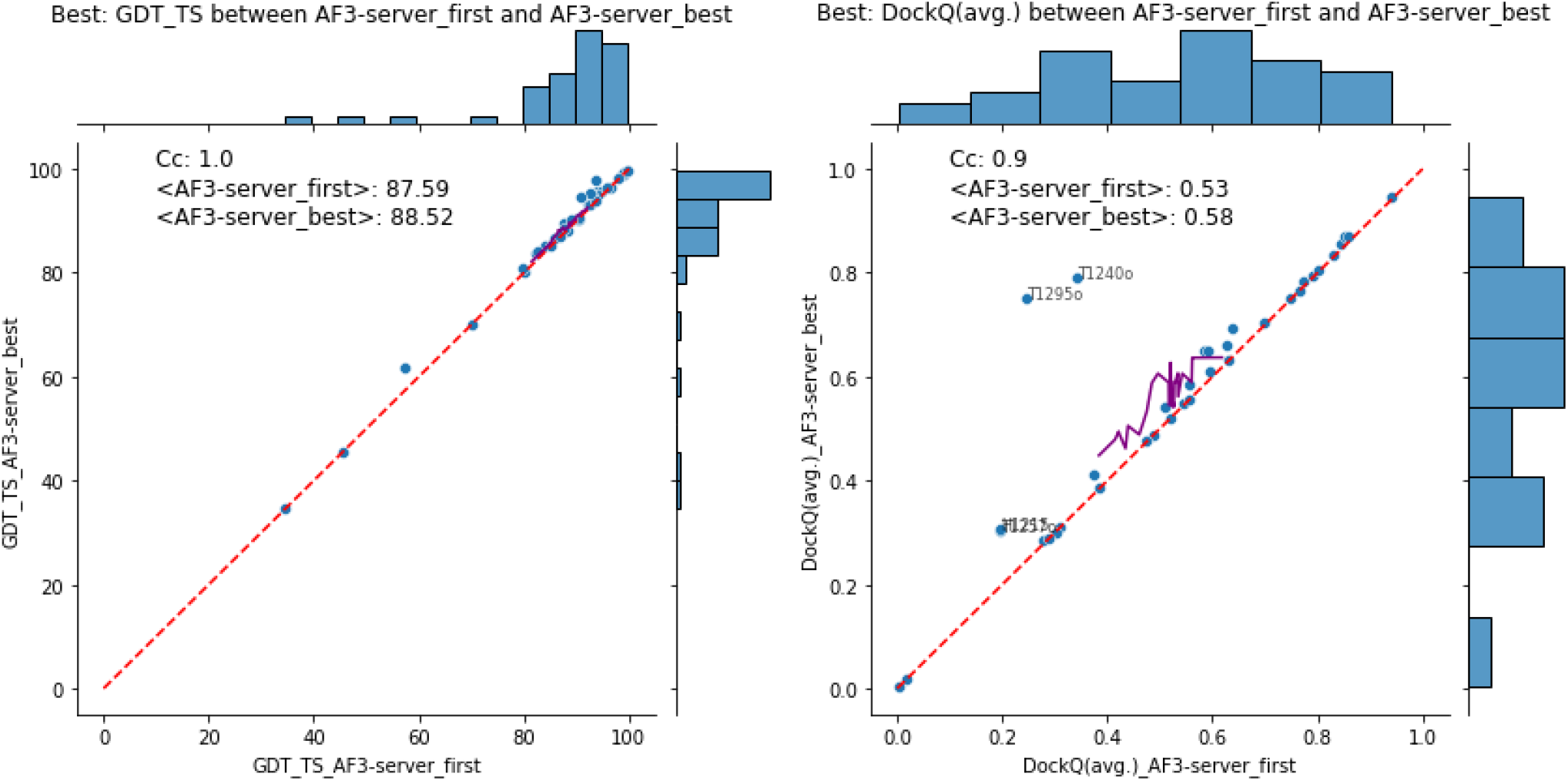
Comparison of the top-ranked versus the best prediction for each target from the AF3-server. The left plot is for domains using GDT_TS, and the right is for complexes using DockQ.

**Figure 6:**
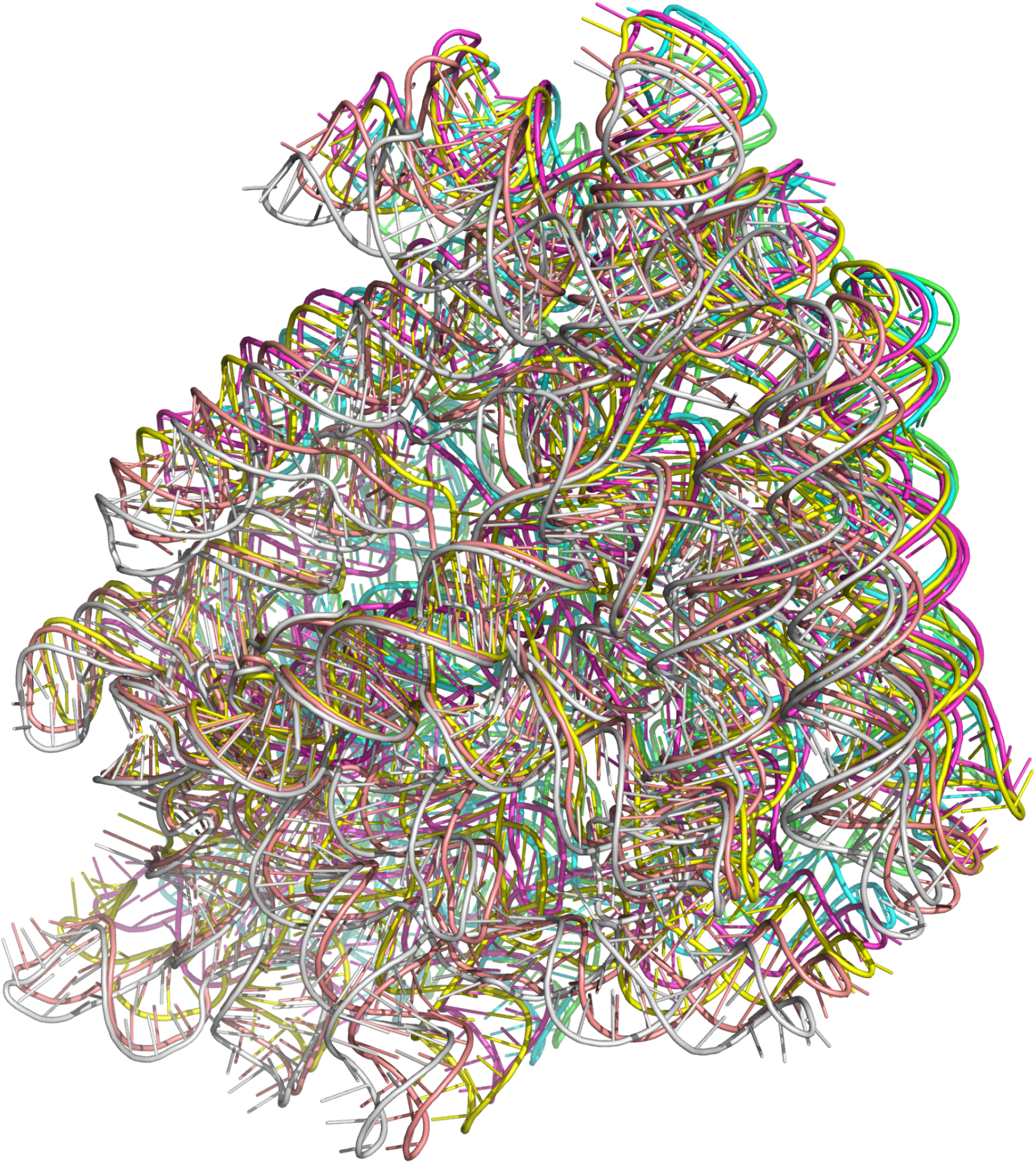
Example of “overlapping” RNA prediction for R1250 (hexamer), each chain is highlighted in one color.

### RNA

In our analysis of AlphaFold3’s performance, we noted that the software frequently predicted overlapping models for multimeric RNA targets, see Figure 7. As this was obvious from visual inspection and the scores, we decided to address this and rerun the predictions until we obtained at least five models without significant overlap and with positive pTM values. In CASP16 official data, the nucleic acid results are categorized into three groups: single RNA, multi-chain RNA, and hybrid complexes. We merged the latter two categories for our analysis since the mixed category contains only a few entries.

**Figure 7:**
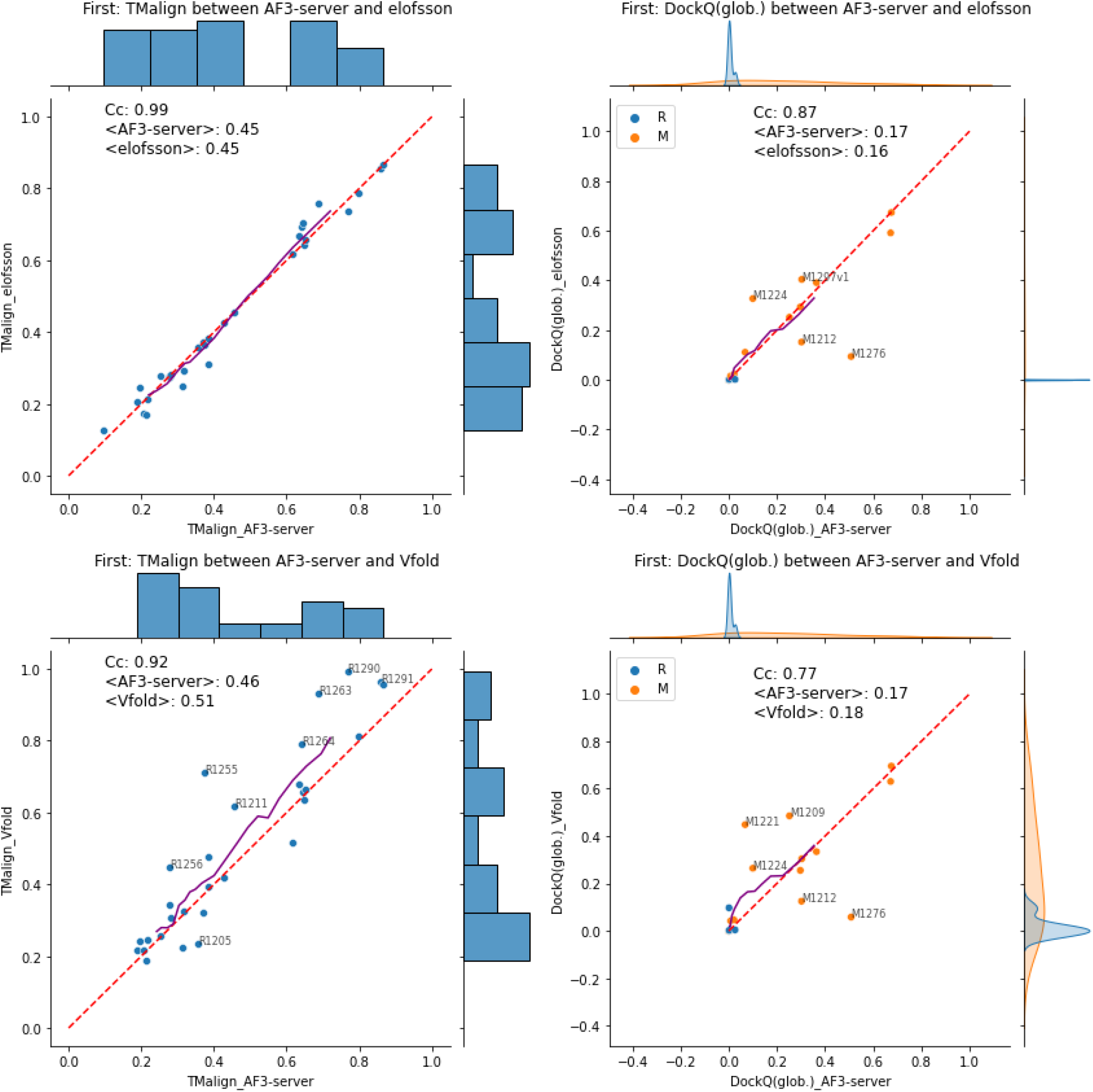
Comparison of RNA predictions. The top row compares the AF3-server with Elofsson predictions, while the second row compares AF3-server with the top-ranked method, Vfold. On the left, the comparison is made at the single RNA level using TMalign, and on the right, the comparison of complexes employs DockQ. Here, the Mixed (M) and pure RNA targets are marked in different colors. Any target with a difference greater than 10% of the maximum value for one of the methods is noted. A rolling average over ten points is shown in purple.

The results show no difference in performance between the Elofsson method and the AlphaFold3 server (see Figure 8). However, it should be noted that there are only a few acceptable models for complexes overall, making direct comparisons difficult. The significance of an increased DockQ from 0.3 to 0.5 can be questioned. Notably, all three methods predicted the same two complexes with a DockQ>0.6, M1293 and M1296, demonstrating consistency across the methods and highlighting that the good DockQ scores stem from accurate predictions of the protein parts of these models. When evaluating single RNA predictions, Vfold slightly outperformed AlphaFold3 on easier targets, particularly when five predictions were allowed. The difference is maintained using the best of five predictions. Here, the average TMscore for single-strand RNAs is 0.55 for Vfold vs 0.48 for AF3-server, and for complexes 0.21 vs 0.19. The difference is minimal for complexes, with some mixed models performing better with one method or the other, although the accuracy, even for the best of the methods, remains low.

**Figure 8:**
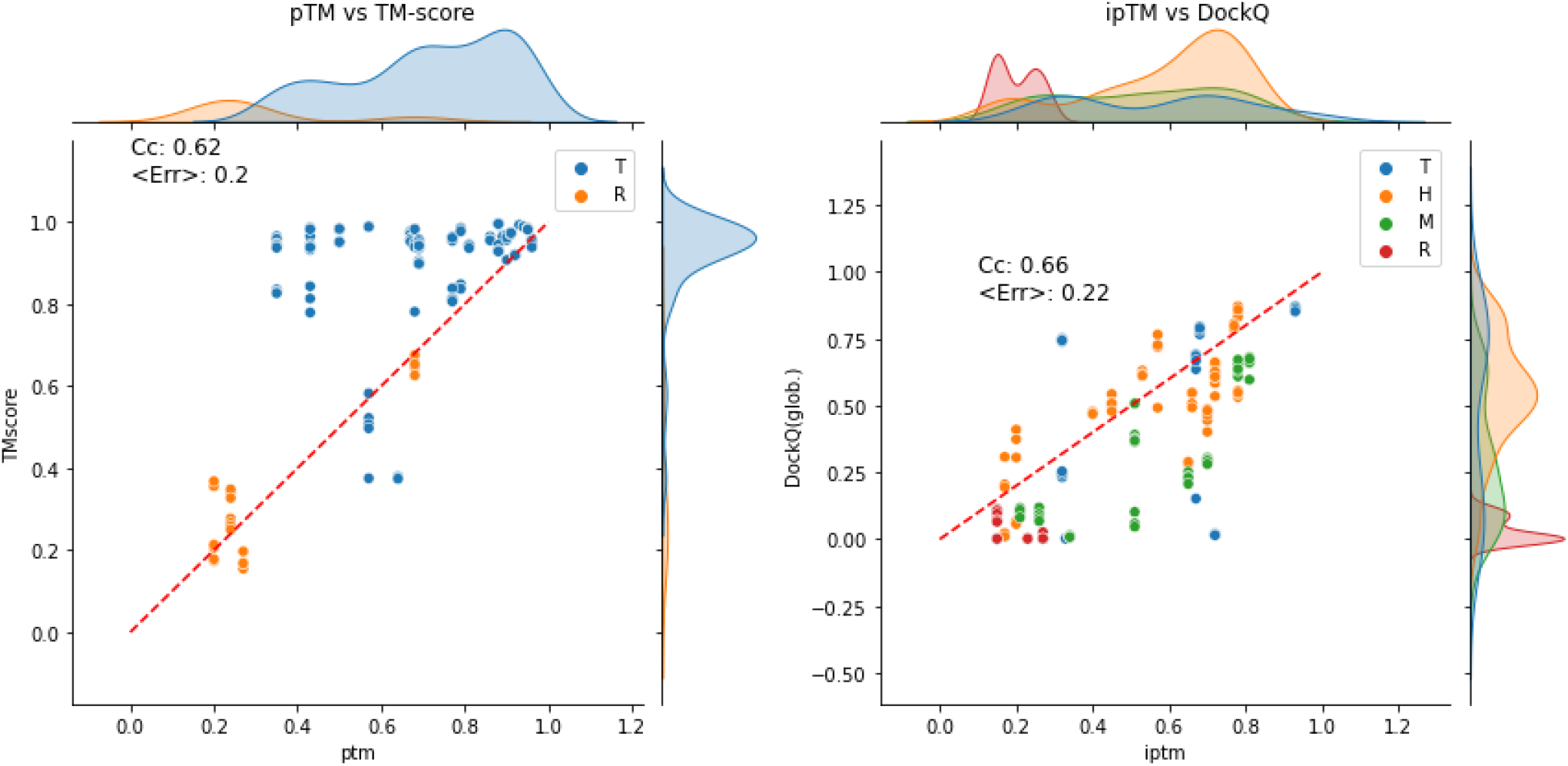
Evaluation of quality estimations. Left plot, pTM vs TMscore, for protein domains and single-stranded RNA. Right ipTM vs DockQ for entire complexes. Each type of complex (T, H, M, and R) is shown in different colors.

### Estimated Model Accuracy (EMA)

When evaluating per-atom Z-score LDDT RMSD, AF3-server and Elofsson were ranked second and third after PLMfold. See the official assessment on the prediction center. Here, we also wanted to compare the predicted TMscores (pTM/ipTM) with the overall quality of a protein/complex. Figure 8 shows that the pTM scores vary significantly between the different protein domains, while almost all domains have a relatively high TMscore, i.e., the correlation is not that good. However, this is caused by the fact that AlphaFold3 predicts the pTM for the entire complex and not for an individual chain/domain. Therefore, the correlation of prediction accuracy is low (Cc=0.66). For single-strand RNA, there seems to be a better agreement of pTM and TM, possibly because these targets were not divided into domains. To provide pTMs for each chain/domain would be a valuable extension to AlphaFold3.

For complexes, we compared ipTM with DockQ, as this estimates the quality of the entire complex. Figure 8 shows that the overall correlation is acceptable (Cc=0.66), but if we compare the pTM with the TMscore, the correlation is better (Cc=0.83), and the average error is 0.13, data not shown. The good correlation can partly be explained by the fact that almost all RNA models are bad, and the pTM scores are also low.

### Stoichiometry

In CASP16, there was a new challenge: to predict the stoichiometry for complexes. We predicted the stoichiometry for 41 stage zero targets, using the AlphaFold3 scores, see Table I. Out of the 41 targets, AlphaFold3 predicted the correct stoichiometry for 14 targets (34%) and 22 (54%) if we include all five submitted models. The Elofsson group did slightly better, correctly predicting one additional first-ranked target and two additional among all submitted targets. The mixed targets (M) were more challenging to submit as both groups only managed to predict one correct stoichiometry for one target (M1287) in the five models submitted. The prediction among all models is better for homomeric (T) targets, but that is likely explained by the fact that there are many fewer options to generate stoichiometries for homomeric targets. Anyhow, it is clear that AlphaFold3 cannot reliably predict the stoichiometry for all targets. Still, at least for the manual predictions for symmetric homomeric targets, it is often possible to guess them correctly using five models.

**Table I:**
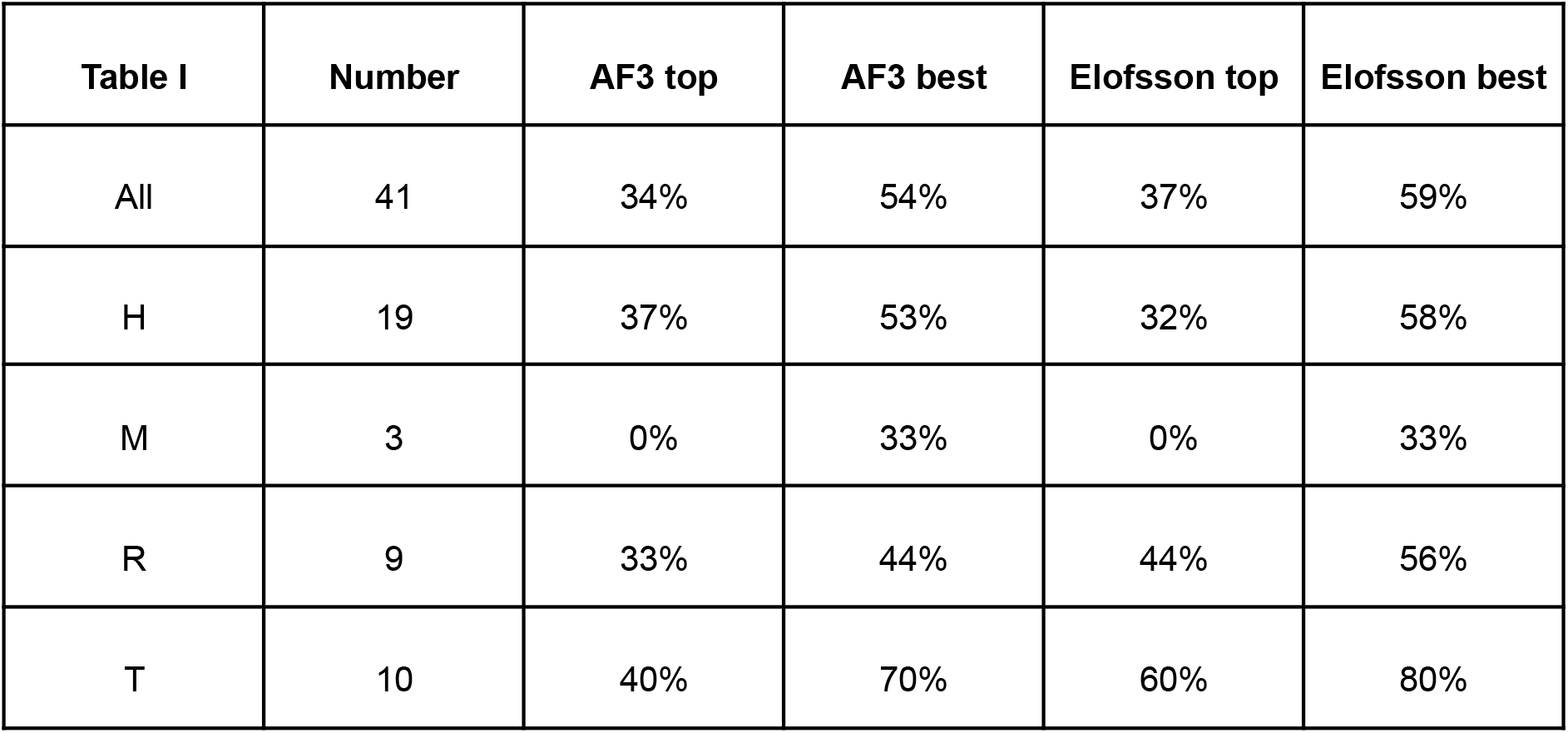
Fraction of correctly predicted stoichiometries, by AF3-server and Elofsson for 41 round zero targets. Results are reported for the top-ranked model or the best of the five submitted models.

## Discussion

There are many ways to evaluate the performance of different groups, and as can be seen above, it is hard to detect the differences using real numbers. Therefore, in CASP, the relative performance, measured by Z-score, is often used to rank the predictions. Using Z-scores highlights minor differences that might have little impact on the real usage of these methods, but help rank the methods. We did not use the Z-scores above, but we assess that they might provide additional information. Therefore, we also compared the ranking of AlphaFold2- and AlphaFold3-based methods using the Z-scores from the prediction center. For simplicity, we decided to use the default evaluation by the prediction center and focused only on the groups’ rank, using the Sum of Z-score>-2.0. Other measures would have provided similar results, but the details would have varied.

For comparison, we also include colabfold_baseline and MassiveFold in the results, as shown in Table II. It can be seen that AlphaFold3 consistently was placed among the top 20% (mainly in the top 10%) of the predictors for All, Easy, and Medium targets, but only in the top 30-40% for the hard targets. Also, it is clear that for all except the hard targets, the predictions are significantly better than for Colabfold_baseline or MassiveFold (both based on AlphaFold2), showing that AlphaFold3 indeed is better than AlphaFold2 for the easier targets. Surprisingly, the performance for both hard single domain targets and hard multimers AlphaFold3 seems not to be any significant improvement over AlphaFold2, and it is clear that massive sampling approaches (see, for instance, the performance of the Wallner group) applied to AlphaFold2 are better than AlphaFold3.

**Table II.** Official rank of the Elofsson (Elo) and AF3-server (Af3) predictors compared with two AlphaFold2-based methods, colabfold_baseline (Co) and MassiveFold (Ma), at the prediction centers website, using the Z-score>-2 measure. The numbers represent the rank of each predictor.

| Table II | All |  |  |  | Easy |  |  |  | Medium |  |  |  | Hard |  |  |  |
| --- | --- | --- | --- | --- | --- | --- | --- | --- | --- | --- | --- | --- | --- | --- | --- | --- |
| Rank | Af3 | Elo | Ma | Co | Af3 | Elo | Ma | Co | Af3 | Elo | Ma | Co | Af3 | Elo | Ma | Co |
| Protein Domains (110 predictors) | 13 | **9** | 40 | 51 | **7** | 9 | 36 | 50 | 19 | **10** | 38 | 54 | 38 | 35 | 50 | **25** |
| Protein Multimers (81 predictors) | **10** | 16 | 30 | 47 | 15 | **8** | 33 | 25 | **2** | 16 | 34 | 39 | 35 | 21 | **20** | 66 |
| RNA/DNA Monomers (64 predictors) | 9 | **6** |  |  |  |  |  |  |  |  |  |  |  |  |  |  |
| RNA/DNA Multimer (32 predictors) | **8** | 12 |  |  |  |  |  |  |  |  |  |  |  |  |  |  |

## Conclusions

We could not detect any improvement in our manual predictions, Elofsson, over the AF3 server. When comparing AlphaFold3 to AlphaFold2, it is noted that AlphaFold3 performs slightly better for complexes; however, this difference becomes negligible when utilizing massive sampling techniques. For domain predictions, no discernible differences were observed between the two versions. Using the official CASP ranking, it was clear that AlphaFold3 performs better on easier targets, but not harder ones. Moreover, while the best methods, which likely incorporate AlphaFold3 as part of their overall pipeline, are only marginally better than the AF3 server for first-ranked models, they show improved performance when selecting the top model out of five submitted, suggesting that in the context of CASP, it remains beneficial to submit a variety of models to enhance ranking. Whether this is useful for usage outside CASP can be discussed, but it highlights that improved selection of the best models would be beneficial. We also show that less than half of the stoichiometries are predicted correctly, indicating an area for future improvement, particularly for non-symmetric complexes. Finally, regarding RNA predictions, it was observed that most models exhibit low quality, with only a slight difference in performance among the best methods and the AF3 server, so RNA predictions remain a challenge.

## Availability

AlphaFold3 was only available as a web server for non-commercial entities during CASP16. Now it is available in source code from https://github.com/google-deepmind/alphafold3 under the “Attribution-Non Commercial-Share Alike 4.0 International” license. All our analysis scripts can be found here: All analysis scripts are freely available from https://gitlab.com/arneelof/CASP16-predictions. Details about all predictions can be found here https://github.com/ElofssonLab/casp16/blob/master/Targets.MD.

## Acknowledgements

AE was funded by the Vetenskapsrådet Grant No. 2021-03979 and the Knut and Alice Wallenberg Foundation. The computations/data handling were enabled by the supercomputing resource Berzelius, provided by the National Supercomputer Centre at Linköping University, the Knut and Alice Wallenberg Foundation, and SNIC, grant Nos. SNIC 2021/5-297 and Berzelius-2021-29.

